# The association between mineralised tissue formation and the mechanical local *in vivo* environment: Time-lapsed quantification of a mouse defect healing model

**DOI:** 10.1101/721365

**Authors:** Duncan C Tourolle né Betts, Esther Wehrle, Graeme R Paul, Gisela A Kuhn, Patrik Christen, Sandra Hofmann, Ralph Müller

**Affiliations:** Institute for Biomechanics, ETH Zurich, Zurich, Switzerland; Department of Biomedical Engineering and Institute for Complex Molecular Systems, Eindhoven University of Technology, Eindhoven, The Netherlands

## Abstract

An improved understanding of how local mechanical stimuli guide the fracture healing process has the potential to enhance clinical treatment of bone injury. Recent preclinical studies of bone defect in animal models have used cross-sectional data to examine this phenomenon indirectly. In this study, a direct time-lapsed imaging approach was used to investigate the local mechanical strains that precede the formation of mineralised tissue at the tissue scale. The goal was to test two hypotheses: 1) the local mechanical signal that precedes the onset of tissue mineralisation is higher in areas which mineralise, and 2) this local mechanical signal is independent of the magnitude of global mechanical loading of the tissue in the defect. Two groups of mice with femoral defects of length 0.85 mm (n=10) and 1.45 mm (n=9) were studied, allowing for distinct distributions of tissue scale strains in the defects. The regeneration and (re)modelling of mineralised tissue was observed weekly using *in vivo* micro-computed tomography (micro-CT), which served as a ground truth for resolving areas of mineralised tissue formation. The mechanical environment was determined using micro-finite element analysis (micro-FE) on baseline images. The formation of mineralised tissue showed strong association with areas of higher mechanical strain (area-under-the-curve: 0.91±0.04, true positive rate: 0.85±0.05) while surface based strains could correctly classify 43% of remodelling events. These findings support our hypotheses by showing a direct association between the local mechanical strains and the formation of mineralised tissue.

## Introduction

It is accepted that organ-scale mechanical loads have a significant influence on the outcome of bone healing. Yet, at the tissue-scale, where healing takes place, the mechanical stimuli guiding this process are unknown. Manipulation of organ-scale loading has been performed via changes in fixation stiffness and defect size, allowing precise application of load to the defect^1,2^. Studies focusing on this manipulation of load have shown differential outcomes^3^. Though understanding of the mechanobiology of bone healing will have profound clinical impact, eventually allowing patient-specific treatment of fractures, a more immediate application of such knowledge might be preclinical research on biomaterials or drugs, via the use of rodent models. In such models, currently a plethora of defect sizes and fixation methods are used^4^. The sensitivity of healing to mechanical stimuli confounds comparisons between studies where: different fixators or defects sizes are used; biomaterials with different stiffnesses are implanted into defects changing the tissue-scale mechanical environment; or pharmaceutical treatments potentially alter the mechanosensitivity of cells. Knowledge of the optimal tissue-scale mechanical conditions would allow compensation for these effects, potentially producing new knowledge on biomaterials and pharmaceutical treatments from existing studies.

At the tissue scale several theories have been presented linking mechanical parameters to tissue differentiation such as deviatoric strain and hydrostatic pressure^5^ or shear strain of the tissue and fluid flow within the tissue^6^. However, these theories cannot be confirmed without spatial and temporal experimental data tracking the healing process^7^. In lieu of this, the mechanoregulatory rules have been implemented within *in silico* models and compared to cross-sectional data of the tissue patterning^8–10^. These state-of-the-art *in silico* models are based upon idealised shapes of the tissue; for example, assuming bone is uniform and cylindrical, and the callus extent is predefined. Unfortunately, histological slices rarely conform to these expectations, impeding quantitative comparison^11^.

An alternative to *in silico* modelling is strain mapping where local deformations are estimated using digital images of tissue in the relaxed and deformed states. This method has been used to parametrise and validate finite element (FE) models^12,13^ and also to correlate tissue phenotypes with mechanical strains^14,15^. While these approaches remove the assumptions used to create finite element models regarding tissue properties, they are sensitive to sample preparation, imaging artefacts and out of plane artefacts given their two dimensionality. Finally, as the data is cross-sectional, the calculated deformation coincides with the tissue compositions but does not precede it.

Time-lapsed *in vivo* micro-computed tomography (micro-CT) produces spatial and temporal experimental data for mineralised tissue and, in the field of bone (re)modelling, has enabled non-destructive examination of (re)modelling events over several weeks^16,17^. When coupled with micro-finite element (micro-FE) analysis, micro-CT imaging, has allowed the investigation of the mechanoregulation of bone (re)modelling^18–20^. To investigate fracture healing, *in vivo* micro-CT has been used to track changes in bone volume (BV)^21,22^. Qualitative associations between strain patterns and bone formation have been observed in two dimensional analysis of time-lapsed micro-CT images and FE simulations^21^.

In this study, we combine time-lapsed *in vivo* micro-CT with micro-FE analysis to quantify the relationship between the local *in vivo* environment (L*iv*E)^23^, bone formation and mineralisation kinetics over the course of healing in a mouse femur defect model. We hypothesize that 1) the tissue-scale mechanical stimulus which precedes the onset of mineralisation is greater in voxels that mineralise than in voxels which do not mineralise and that 2) this phenomenon is independent of the magnitude of global mechanical loading of the defect.

Additionally, reconciling the wide range of densities that are observed during the fracture healing process is key to improve micro-FE prediction of mechanical stimuli. Traditionally, either single absolute thresholds, ranging from 394.8 to 641 mg HA/cm^3^ ^24–26^ (HA: hydroxyapatite); or relative thresholds based on percentages of grey values; such as 25% to 33%^27–29^ have been used to segment bone. We improve upon these approaches via the use of a “multi-density threshold approach”, whereby we apply a range of thresholds to identify the spatially and temporally changing densities of bone, allowing us to quantify local mineralisation kinetics and reconcile the range of bone densities within the healing environment.

## Results

### Longitudinal monitoring of fracture healing

Defect healing followed a typical pattern, progressing from the reparative phase to bridging and then displaying mineralisation and remodelling. The 0.85 mm defect healed in 9/10 mice with the single atrophic non-union which was excluded from analysis. The 1.45 mm defect group had a non-union rate of 4/9, similar to the results of Zwingenberger et al.^30^. The 0.85 mm group qualitatively had a larger callus compared to the 1.45 mm group (Figure 1). As mentioned before, there is no consensus in literature regarding the level of mineralization for segmenting bone during the healing process. The apparent progression of healing was shown to be highly sensitive to the threshold in both groups. When comparing the highest threshold to the lowest, the peak BV/TV was delayed by at least a week for all regions (Figure 2 a-h). As seen in the formation and resorption rates, two phases of mineralisation could be distinguished (Figure 2 i-p). Firstly, an initial deposition of a large amount of low-density tissue occurred in week 2 and 3, followed by a production decay from the third week onwards; and secondly, an increase in the resorption of low-density tissue and a peak in the formation of high density tissue directly following the first phase (Figure 2 i-p). These results indicate that the production of a lowly mineralized callus and maturation of the tissue are independent processes. The resorption of bone in the FC volumes (Figure 2 k&o) also confirms the results of Schell et al.^31^ who observed osteoclastic activity early in the healing process. Our results indicate that in all VOIs resorption can occur a week after material is deposited, highlighting the substantial overlap between the reparative and remodelling phases.

**Figure 1.**
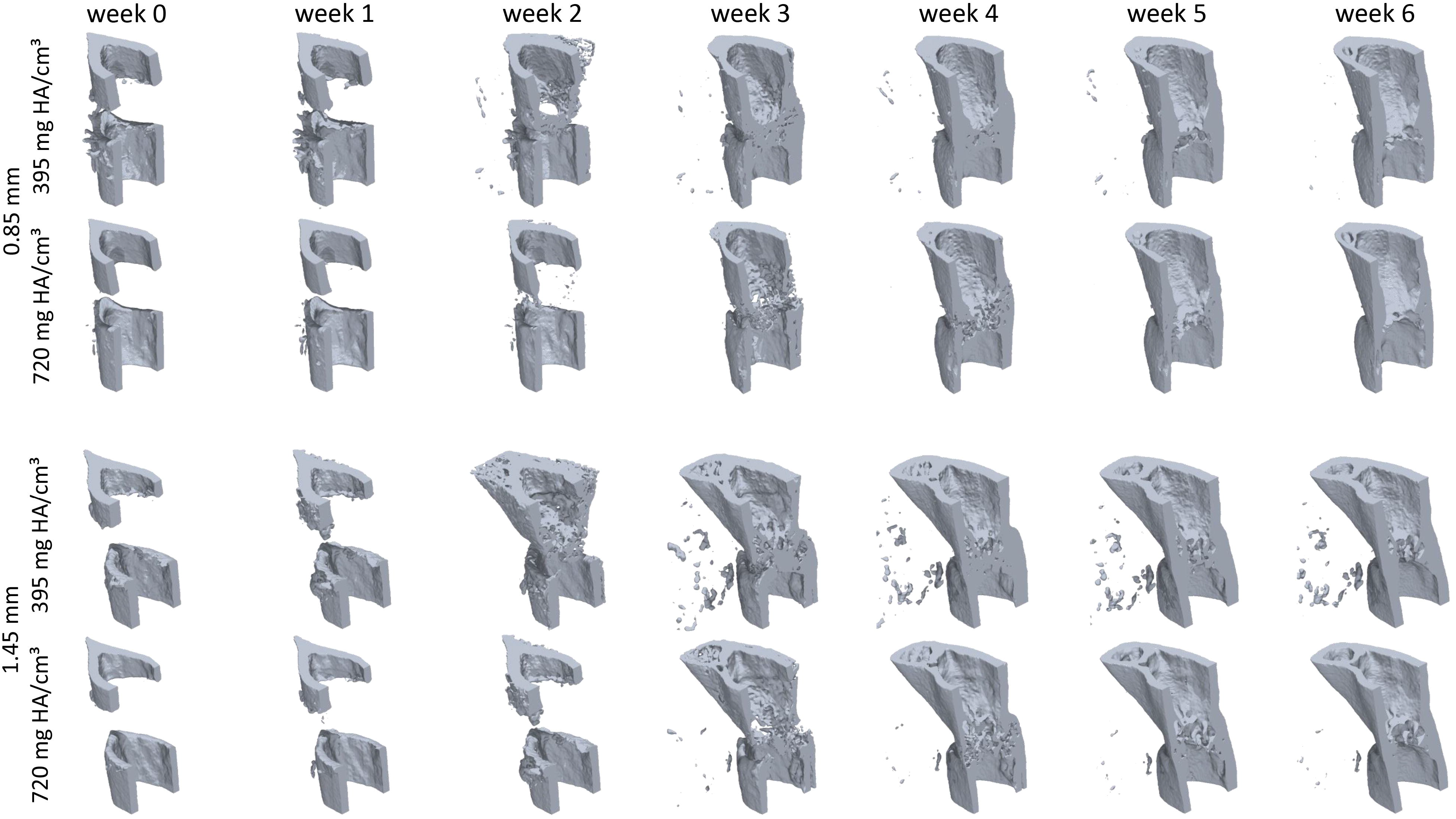
Callus development over the course of the study for both 0.85 mm and 1.45 mm defects. Two thresholds are shown for each mouse, most significant is the difference between observed state of healing at week 2, where for the lower threshold both groups are fused and for the higher threshold no healing has occurred.

**Figure 2.**
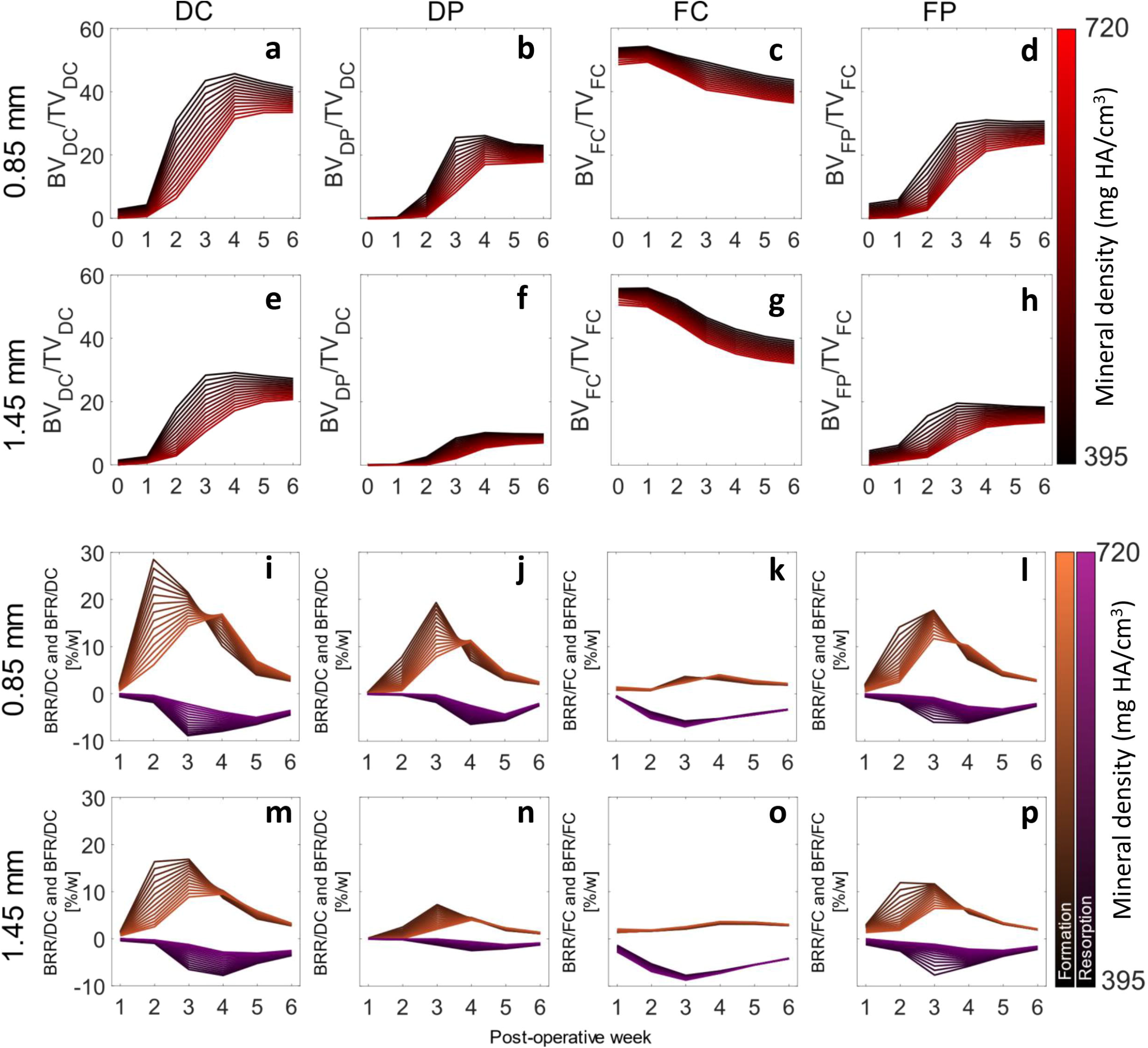
The development and kinetics of mineralised tissue in the defect. a-h show the time course of bone volume. i-p shows the mineralisation kinetics, resorption and formation are separated. The mean of the 0.85mm, panels a-d and i-l, and mean of 1.45 mm, panels e-f and m-f. Note that as a proportion of the defect more tissue is formed within the 0.85mm group compared to the 1.45mm group. A successful healing outcome is a union in which the bone is indistinguishable from the original cortical bone, comparing the original cortical fragments FC+FP and defect VOIs DC+DP between groups, it is apparent that the 0.85mm defect has similar BV/TV in the DC+DP (a+b) and FC+FP (c+d) in the final weeks, while the 1.45mm defect groups has lower BV/TV within the DC and DP regions. This shows that insufficient bone is produced in the larger defect. For both groups the difference between the highest and lowest threshold would cause relative underestimation of callus size in the initial 3 post-operative weeks. The rates of mineralisation were threshold dependent, with an initial large deposit of lowly mineralised tissue followed by a second phase of maturation a-h. Resorption of the existing cortical fragments was apparent in week 1-3.

### Estimation of physiological loading

To determine physiological boundary conditions, a load estimation algorithm was used on intact femoral midshafts from a second cohort of mice. The load estimation algorithm predicted an axial compressive load of 10.0 ± 1.9 N with a bending moment of 3.5 ± 0.7 Nmm.

### Mechanobiology of tissue mineralisation in the defect

The associations between the effective strain and the mineralisation events in the defect volume were strongest in the second post-operative week, this was true for both groups (Figure 3 a&d). This corresponded with the initiation of hard callus formation (Figure 1 & Figure 2). In both cases, the area under the curve was reduced for the third week and this decrease was proportional to the threshold value. The higher density bone in the third week consisted of both new bone and the maturation of low-density bone deposited in week 2. An AUC of 0.5 implies that maturation was not associated with the mechanical stimuli. This trend is reversed in later weeks, indicating that bone deposited during the remodelling phase is again under mechanical control. For the 0.85 mm group, an effective strain of greater than 0.51±0.25 was found to provoke bone formation, while for the 1.45 mm defect the effective strain was 0.85±0.34. Excluding mice in which non-unions occurred, that effective strain was 0.54±0.26. While not significant, this may indicate that the non-union mice might have cells that are less mechanically sensitive.

**Figure 3.**
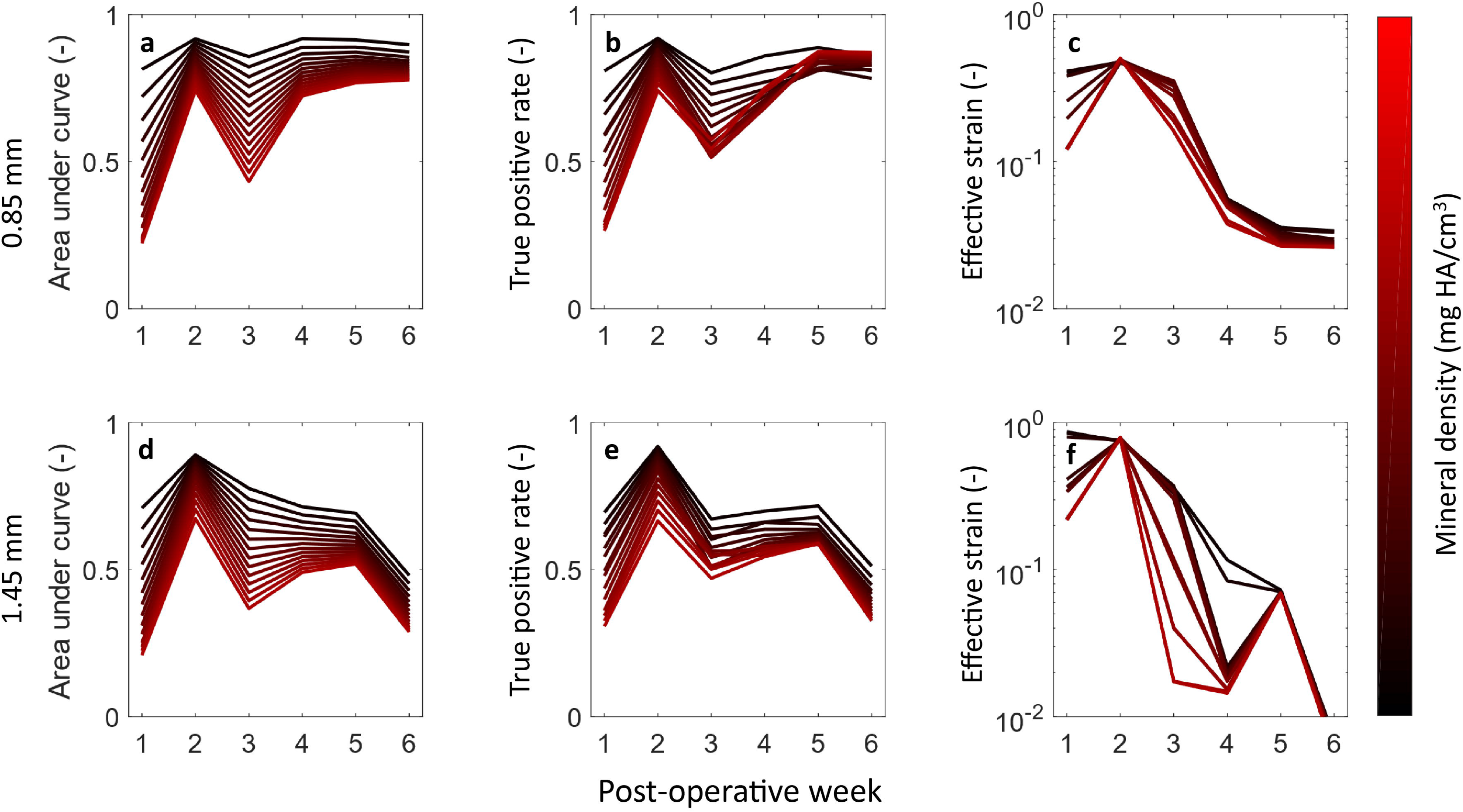
The associations between mineralisation on the mechanical local *in vivo* environment: Time course of area under curve (AUC) (a,d), True positive rate (b,e) and Effective Strain thresholds (c,f) for 0.85 mm (upper row) and 1.45 mm (lower row) defects. a,d) The area under curve for the 0.85 mm and 1.45 mm respectively. In both groups the AUC was maximum in the second week. The AUC declined in the third week for both groups, this decline was largest for the highest mineralised tissue, again recovering for the 0.85 mm, while continuing to decline for the 1.45 mm group. The poor association between the formation of highly mineralised tissue and mechanical signal in the third week is an indication that the maturation of bone is independent of the local mechanics while its formation is dependent. The recovery of the AUC in the later weeks for the 0.85 mm is likely due to the group containing only mice with unions. The true positive rate of the optimal threshold (b,e), initially show a similar trend to the AUC. However, for the 0.85 mm group the association for the highest mineral density threshold is also highest in the final week, indicating that controlled remodelling is taking place. The optimal effective strain thresholds (c,f) show a similar level of strain in the second week. A large amount of variation with respect to mineral density threshold can be seen in the 1.45 mm group from week 3 onwards, which is likely due to the presence of both unions (which stress shield soft tissue) and non-unions in the group.

The reduction in the effective strain which provoked mineralisation over time (Figure 1, c&f) represents the stress shielding caused by newly formed bone. The effective strain for the 1.45 mm group was lower in week 6 due to the large number of non-unions, which had significantly less strain in the bone tissue.

### Mechanobiology of the surface

In order to associate the (re)modelling events on the bone surface with the mechanical stimuli on the surface, the bone surface of the preceding week was extracted from the overlaid binarized images. The state of the surface was recorded, i.e. whether it was a site of formation, resorption or quiescence. The mechanical stimuli were calculated using micro-FE and extracted at the corresponding surface locations. Two thresholds (resorption and formation) were then used to predict the surface state which was then compared to the *in vivo* derived ground truth. Heat plots were generated for each week, mineral density threshold, and VOIs. Only the lower right triangle is occupied as the threshold for resorption and formation cannot overlap. This analysis was performed for all mice in the 0.85 mm group which satisfied the exclusion criteria, using all thresholds and time-points (Figure 5d). In the defect, the CCR in the first 3 weeks was approximately 35-37%, this rose to 40-43% by the 6^th^ week (Figure 4b). In the VOIs of the cortical fragments, the CCR was initially 50-56% decreasing to 39-40% by the 6^th^ week (Figure 4a). At lower mineral density thresholds, for both the defect and cortical fragments, there was a tendency for a higher CCR. Comparing the defect and cortical fragments, it is apparent that the CCRs for both DF+DP and FC+FP approach the same value in the final week (Figure 4).

**Figure 4.**
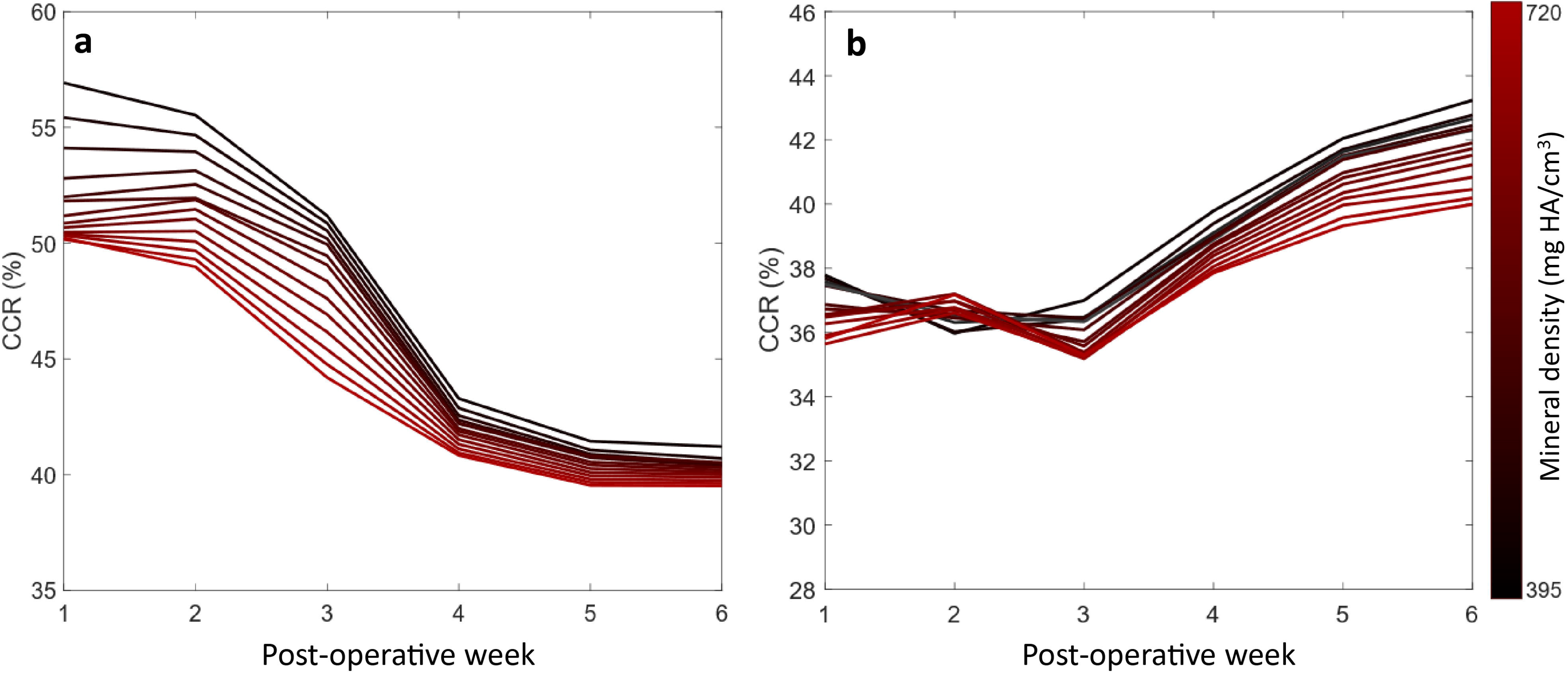
The highest mean correct classification rates for remodelling events for each week and mineralisation threshold. a) Remodelling events in the fragment VOIs, showing a drop in CCR as the cortices shift from resorption to balanced remodelling. b) Remodelling events in the defect, initially close to random followed by an increase each week post bridging.

## Discussion

The aim of this study was to investigate the mechanical regulation of bone healing in the local *in vivo* environment (L*iv*E) and determine if the tissue-scale stimulus preceding formation of mineralised tissue was independent of global loading of the defect. Our first hypothesis, that the formation of mineralised tissue during bone healing is regulated, at least partly, by tissue scale mechanical stimuli could be confirmed, with up to 91±3.4% of mineralised tissue formation within the defect being predicted by a single threshold for both defect sizes (Figure 3). The lack of a significant difference between the optimal *ε_eff_* during the 2^nd^ week post-operative measurement of the 0.85mm and 1.45mm defect groups further strengthens this finding and means that we cannot reject the second hypothesis.

While previous studies have focused on qualitative assessment of the mechanoregulation^21^, or on two dimensional cross-sectional data^5,10,15^, this study uses fully three dimensional information and associates a preceding strain with the tissue. In our study, we focused on *ε_eff_* as a classifier. The choice of *ε_eff_* rather than SED is due to the heterogenous nature of the tissues in the healing bone. As SED scales linearly with stiffness, this precluded comparison between regions with different Young’s moduli. In contrast, *ε_eff_* is independent of the element elasticity. Previous studies have attempted to divide the mechanical stimuli into either shear and volumetric components or the perfusion of fluid through the tissue matrix and shear^5,6^. However, adversarial testing by Repp et al.^10^ could not falsify any proposed theory. Nevertheless, Repp et al. found that a simple model using only volumetric strain could produce an equivalent healing outcome. In contrast, Epari et al.^32^ stated that deviatoric strains dominated the mechanoregulation in their studies. The difference between Epari et al. and Repp at al. might be explained by the different dominant modes of loading: shear versus compression. Our use of *ε_eff_* combines both deviatoric strains and volumetric strains into a single scalar quantity. As SED has been used extensively for the analysis of bone (re)modelling, using a derived quantity can allow for a unifying theory of both bone repair and (re)modelling.

When considering (re)modelling, the association of (re)modelling events at the surface is comparable to Schulte et al.^33^, who were able to predict the spatial accuracy of formation and resorption (equivalent to CCR) at 47.6 % (SD=3.3 %) in caudal vertebrae of mice compared to the CCR of 42% (SD=2.4 %) in this study. The surface (re)modelling events of the cortical bone initially had a strong association with the mechanical stimuli when compared to bone in the defect. The CCR of cortical bone decreased over the study period, converging with the CCR of the defect (Figure 4). We see two potential explanations for this behaviour: Firstly, the cortical VOIs were inactive during the first week (Figure 2 k&l). Thus, most of the surface was in a quiescent state with a small amount of bone resorption. This reduced the total amount of possible misclassifications as the ground truth has two states rather than three. The amount of (re)modelling activity then increased in the subsequent weeks, hence the CCR decreased as the surface enters a state of balanced remodelling where the amount of newly formed bone is equal to the resorbed bone. Secondly, it is possible that errors in our boundary conditions were lower for the non-healed bone when the majority of load was transmitted through the external fixator and boundary artefacts relating to the screws were lower. In the coarse FE-model, the entire callus is modelled as a homogenous material where the boundary conditions are related to the *in vivo* micro-FE model based purely upon the axial stiffness of the callus. The bending stiffness of the callus is not controlled in either model and is a source of error.

There were several limitations related to this work. The following assumptions were made in our FE models regarding material properties: (i) all materials are linear elastic, (ii) all materials have a homogenous Poisson’s ration of 0.3, (iii) all voxels not containing bone have a homogenous stiffness of 3 MPa, and (iv) the boundary conditions are estimated. Regarding limitations (i) and (ii), the first two assumptions are constrained by model size. The largest model had approximately 102 million degrees of freedom and to solve these models in a reasonable amount of time and have a manageable amount of data, we relied upon the linear elastic micro-FE solver ParOSol^34^. ParOSol has the restriction that all elements have the same Poisson’s ratio. Steiner at al.^13^ used electro speckle interferometry to strain map sections of calluses to assess the Poisson’s ratio of tissues in a healing callus, and while they found high variance in the locally assessed Poisson’s ratio, the average Poisson’s ratios of cartilage and bone were 0.3, indicating that our assumption is in line with the literature. Regarding assumption (iii), using our micro-CT imaging protocol, it was only possible to differentiate mineralised tissue from non-mineralised tissue. Therefore, the development of soft tissues and their association with the mechanical stimulus was not quantified. Incorporation of MRI data for comparable fixator stiffness and defect size, such as that captured by Haffner-Luntzer et al.^35^, would allow our analysis to include cartilage formation and more detailed mechanical stimulus in the soft tissue region. Finally considering assumption (iv), the physiological loading of the mouse hind limb is currently unknown. Although a musculoskeletal model of the mouse hind limb has been developed by Charles et al.^36^, this has not been validated or applied to mouse gait data. Charles et al. calculated the maximum moments exerted on a mouse femur. The muscle M. psoas major induces a peak flexion moment of 3 Nmm on the femur. This muscle is inserted into the femur at approximately the same level as the boundary of the micro-FE model used for load estimation. This moment is comparable to the 3.5 ±0.7 Nmm we calculated using a load estimation algorithm. The prediction of such similar values with two different methodologies lends confidence to the predicted physiological loading presented in this paper.

In summary, we investigated the association between formation of mineralised tissue and the local mechanical stimuli over the course of healing in a mouse femoral defect model. The results indicate that the bone healing process is mechanically regulated at the tissue-scale in the reparative phase. These results also provide both parameters and ground truth to improve and validate fracture healing models in mice. Combinations of the methods presented will reduce animal numbers needed in bone healing studies, through longitudinal monitoring, and allow quantification of mineralisation kinetics and mechanosensitivity. Finally, knowledge of mechanical stimuli at the tissue-scale will allow mouse studies to choose the fixation stiffness and defect sizes which tailor the local mechanical environments to physiologically relevant ranges.

## Material and Methods

### Animal model and imaging

All animal procedures were approved by the authorities (licence number: 36/2014; Kantonales Veterinäramt Zürich, Zurich, Switzerland). We confirm that all methods were carried out in accordance with relevant guidelines and regulations (ARRIVE guidelines and Swiss Animal Welfare Act and Ordinance (TSchG, TSchV)). The study comprised of two groups of female C57BL/6J mice, age 20±1 week (Janvier Laboratories, Le Genest-Saint-Isle, France). In both groups, a femoral defect was created by first stabilising the femur with a radio-translucent external fixator (MouseExFix, RISystem AG, Davos, Switzerland) and then removing a section of bone. One group (n=10) received a 0.85±0.09 mm defect. The second group (n=9) consisted of animals from an existing study^37^, which had a 1.45±0.16 mm defect. The mice were scanned weekly using *in vivo* micro-CT (vivaCT 40, Scanco Medical AG, Brüttisellen, Switzerland) with an energy of 55 kVp, an integration time of 350 ms and a current of 145 uA with 500 projections. The scanning period was 6 weeks, totalling 7 measurements, including a post-operative scan. To prevent motion artefacts during scanning, the external fixator was secured in a custom designed holder. During all experimental procedures, the animals underwent isoflurane anaesthesia (Induction: 5%, maintenance 2-3%). Analgesia (Tramadol, 25 mg/l; Tramal^®^, Gruenenthal GmbH, Aachen, Germany) was provided via the drinking water during the peri-operative period (two days before surgery until the third postoperative day). Atrophic non-unions, qualitatively assed in the micro-CT scans, served as an exclusion criterion.

### Image processing

Three dimensional micro-CT images were reconstructed at an isotropic nominal resolution of 10.5 μm. Images were registered sequentially; proximal and distal bone fragments of unbridged defects were registered separately as it was found that small relative movements between proximal and distal bone fragments occurred between measurements, the registration used the algorithm described by Schulte et al.^17^. After image registration, the images were Gaussian filtered (sigma 1.2, support 1) to reduce noise. The multi-density threshold approach was then applied, whereby images were binarized with thresholds ranging from 395 to 720 mg HA/cm^3^ in steps of 25 mg HA/cm^3^. The threshold of 395 mg HA/cm^3^ corresponded to the lowest value used to segment bone in existing studies^25^. The highest threshold was chosen as it allowed the threshold range to encompass 640 mg HA/cm^3^, the highest value found in the literature^24^.

The bone volume was evaluated in four non-overlapping volumes of interest (VOIs) which were created from the post-operative measurement. These consisted of a defect centre (DC), defect periphery (DP), a volume encompassing each both cortical fragments (FC), and the fragment periphery (FP) (Figure 5a). These volumes were identified automatically using the following algorithm: The input was the post-operative image binarized with a threshold of 645 mg HA/cm^3^, which was chosen because it disconnected surgical debris from the cortical fragments. The cortical fragments were found using component labelling. The marrow cavities on each side were flood filled, creating the FC VOI. The cortical surface was taken from the last filled slice of the FC mask and interpolated linearly across the defect. A raytracing approach was implemented, where rays were cast from the image boundaries perpendicular to the axis joining the centre of mass of the cortical fragments. If the ray struck a cortical fragment (Figure 5a) all voxels along its path were included in the FP VOI, while if the ray struck the interpolated surface the voxels were then added to the DP VOI (Figure 5a). The DC VOI was then determined as the voxel in the image not included in the FC, FP and DP VOIs. The bone volume was quantified for all thresholds. The volumes were normalised to the central VOIs, DC for DP and FC for FP representing the total volume (TV) of intact bone, which can be considered as a target for healing.

**Figure 5.**
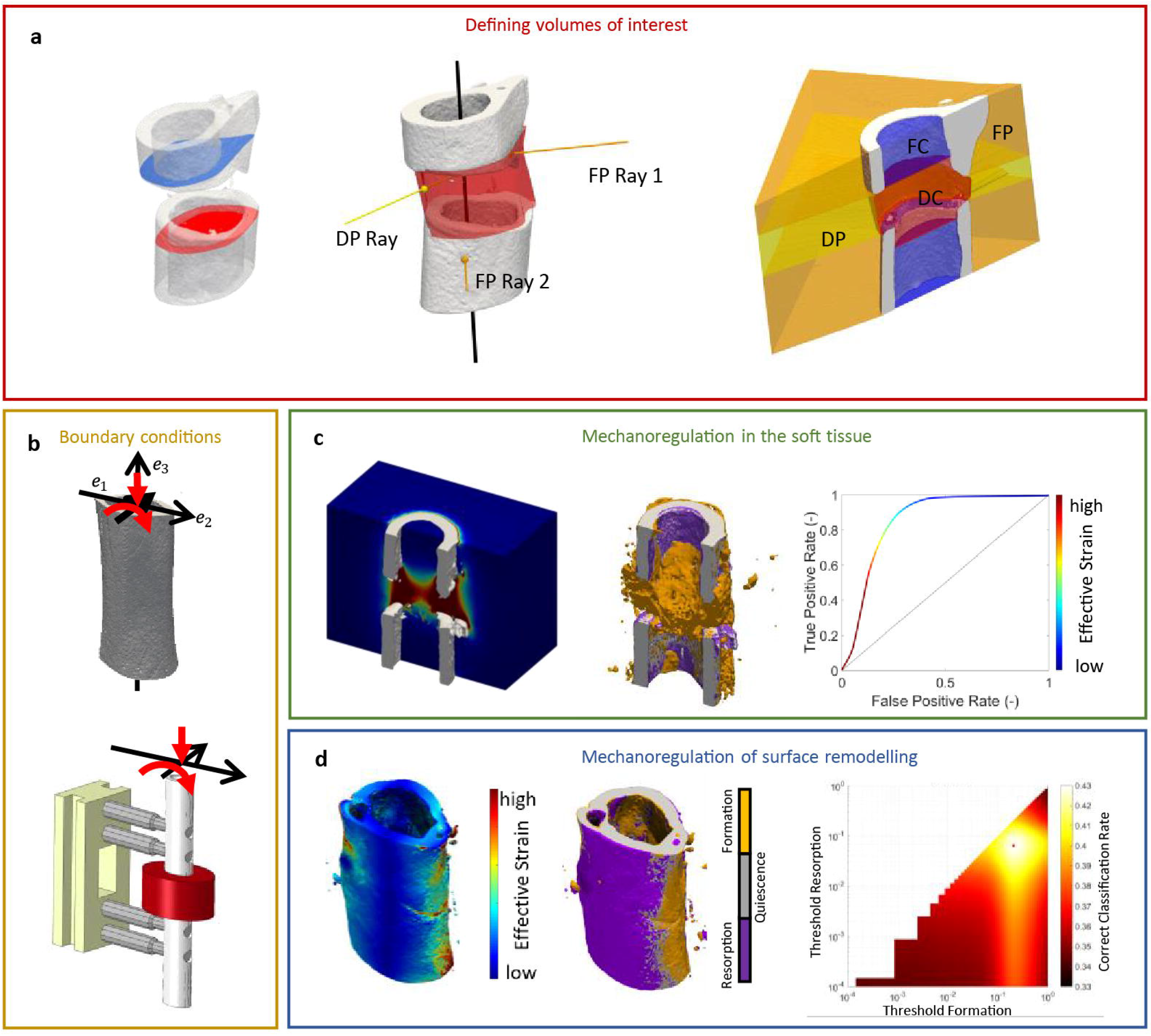
a) Illustration of ROI definition process: First the first and last intact cortical slice in the imaging plane are found (proximal blue, distal red). These shapes are then interpolated across the defect, providing an estimation of the original periosteum. Rays are then cast from the image boundaries along normals to the major axis of the fragments. Rays which intersect the cortical fragments are classified as being members of FP, while rays which intersects only virtual periosteal surface (red) are classified as being in DP. The DC volume is then determined as the space not occupied by FC, FP or DP. b) The pipeline for FE analysis: initially the load on the femur was determined using load estimation on intact contralateral femurs. These loads were then applied to a coarse FE-model which determined the boundary conditions for the *in vivo* micro-CT model. c) The association between mineralised tissue formation and mechanical strain in the soft tissue, micro-FE analysis is used to calculate the strains in the soft tissue, using the time-lapsed *in vivo* micro-CT images as ground truth ROC analysis is performed determining how effective mechanical strain is as a predictor. d) Association of remodelling events with mechanical strain. Using the micro-FE results for the mineralised tissue a set of thresholds are swept through the strain space classifying formation and resorption events. Using the time-lapsed *in vivo* micro-CT images as ground truth, a correct classification rate for each set of thresholds is determined.

### Finite element analysis

The mechanical signal was calculated using three hierarchical FE analyses; 1) Load estimation was used to determine the physiological loading on the intact mouse femur (Figure 5b), 2) A coarse finite element model of the bone-fixator system was used to determine the boundary conditions of the femoral defect (Figure 5b), and 3) a micro-FE analysis based upon the *in vivo* micro-CT images was used to estimate the tissue-scale mechanical stimuli (Figure 5c&d).

### Load estimation

The physiological loading of the mouse femur was determined using a load estimation method proposed by Christen et al.^38^. This method uses the underlying microstructure of the bone to determine a combination of load cases, which create a homogenous strain distribution at a physiological tissue loading level.

The loading parameters were estimated from images of intact contralateral femurs of a second cohort of seven female mice (C57BL/6J) from a previously published fracture healing study^39^. Samples were scanned and processed *ex vivo* in a desktop micro-CT system (microCT 40, Scanco Medical AG, Brüttisellen, Switzerland) according to the protocol established by Kohler et al. ^40^. The mid-shaft was extracted manually as the region above the lower growth plate and below the lesser trochanter. Two load cases with unitary loads were applied to the mid-shafts: axial compression and a bending moment around the minor axis of ellipticity of the femoral cross-section (Fig 5b). The micro-FE analyses for each load condition were then solved using ParOSol^34^, using a 1:1 conversion of voxels to elements. The load estimation algorithm was then used to determine the physiological loading applied to the shaft.

### Coarse finite element model

To translate the organ-scale loads to the tissue-scale in the *in vivo* micro-CT model, a coarse finite element model was created containing the external fixator and the femoral mid-shaft. The central section of the mid-shaft was replaced with a material representative of the *in vivo* micro-CT model. The model was solved using ABAQUS V6.11 (Dassualt systems, Vélizy-Villacoublay, France). The boundary conditions consisted of the load and bending moment determined using the load estimation. All elements were linear elastic, bone stiffness was 14.8 GPa and a Poisson’s ratio of 0.3 was used ^41^. The femoral mid-shaft was modelled as elliptical tubes, with a thickness of 0.2 mm and a minor and major axis of 1.0 mm and 2.0 mm respectively. The fixator was modelled on the MouseExFix system used in the study. The model of the entire system consisted of approximately 250,000 elements. The validity of this model was verified through comparing the simulated stiffness to mechanical compression experiments of the external fixator implanted into PMMA cylinders with an empty defect (Supplementary Fig. S1 and Supplementary Table S1).

### Estimation of *in vivo* strains

For each mouse and time-point a micro-FE model was created. Each model combined the thresholded images, where HA-equivalent mineral densities were converted to a Young’s modulus using a linear relationship^42,43^ with a value of 14 GPa corresponding to 720 mg HA/cm^3^, the highest level of density segmented, while 395 mg HA/cm3 corresponded to 4 GPa. The background of the image was given a value of 3.0 MPa to represent soft tissue^44^. The stiffness of the defect was determined by applying a simple compression of 1% displacement to the cortical bone and the marrow cavity. Using the coarse model, physiological displacements at the image edges could be determined and applied as boundary conditions. The mechanical stimulus used was effective strain was calculated as described by Pistoia et al.^45^. The micro-FE models consisted of approximately 25 million elements, totalling 75 million degrees of freedom, which were solved with the parallel solver ParOSol^34^. Simulations were run on Piz Daint, a Cray XC30/40 system at the Swiss National Centre (CSCS), utilising 8 nodes and 144 cores with a solution time of approximately 5 minutes per analysis.

### Analysis of mechanobiology of bone healing

To associate bone formation with the mechanical stimuli within the defect, a receiver operating characteristic analysis (ROC) was applied (Figure 5c). In our application of ROC, the ground truth was created by overlaying binarized images where newly mineralised bone was identified as condition positive, while tissue which did not mineralise was condition negative. The ROC curve was created via sweeping a threshold through the *ε_eff_* space, values above the strain threshold were classified as bone while below were considered soft tissue. The comparison to the ground truth created the ROC curve (Figure 5c). The analysis was performed in the DC and DP volumes. This process was repeated for each mouse, time-point and mineral density threshold. Due to the large amount of data, the analysis was summarised in terms of area under the curve and the true positive rate selected was the furthest from the random classification line at 45°.

### Analysis of mechanobiology of callus remodelling

On the bone surface, the prediction of sites of formation, resorption and quiescence using the preceding effective strain was interpreted as a multi-class classification problem. The classifier function *f_j_* consisted of two thresholds, as shown in equation (1). The lower threshold classified the sites of resorption (*T_R_*), and an upper threshold classified the sites of formation (*T_F_*), values between the thresholds were classified as quiescent.

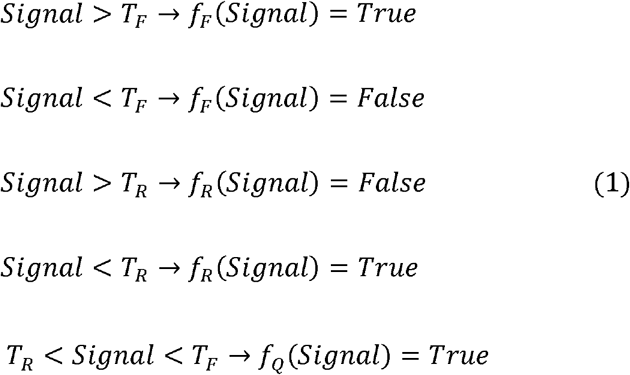

This classifier corresponds to the simplest possible mechanostat model, having just two parameters (*T_R_ and T_F_*)^46^.

The bone surface was defined as the interface between the bone and the background using a 3D ‘von Neumann’ neighbourhood with a radius of 1 voxel^47^. The ground truth *G* was determined with sites of formation defined as the interface between quiescent and newly formed bone. Sites of resorption were defined as the interface between resorbed bone and the background, while the quiescent surface was the interface between quiescent bone and the background. The predicted states were then compared to the ground truth and a category-wise normalised confusion matrix *C*, equation (2), was generated.

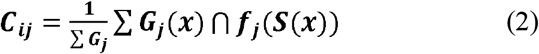

Diagonal entries are analogous with true positive rate, while off diagonals are false negatives and positives. The trace of the normalised confusion matrix indicates the overall performance for all categories. The normalisation step compensates for differences in the number of surface voxels in each category (formation, quiescence and resorption), and thus gives all events equal weighting. An exhaustive analysis comparable to ROC was performed in which both thresholds were swept through the strain space generating two dimensional heat charts (Fig 5d). The colour represents the trace of the confusion matrix divided by three to give an average correct classification rate (CCR), as described in equation (3).

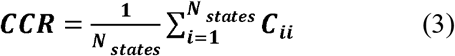

This analysis was performed for each mineral density threshold level, and the maximum CCR was determined for each image and time-point.

### Data availability

All necessary data generated or analyzed during the present study are included in this published article and its Supplementary Information files (preprint available on BioRxiv/2019/721365). Additional information related to this paper may be requested from the authors.

## Acknowledgements

The authors gratefully acknowledge funding from the European Union (BIODESIGN FP7-NMP-2012-262948) and computational time from the Swiss National Supercomputing Centre (CSCS). E. Wehrle received funding from the ETH Postdoctoral Fellowship Program (MSCA-COFUND, FEL-25_15-1).

## Author contributions

The study was designed by D.C.T, E.W, G.A.K, S.H and R.M. The experiments were performed by E.W, G.R.P and G.A.K. Data analyses were performed by D.C.T. Interpretation of the data was performed by D.C.T, E.W, G.R.P, G.A.K, P.C and R.M. The manuscript was written by D.C.T and reviewed and approved by all authors.

## References

1. Goodship, A., Watkins, P., Rigby, H. & Kenwright, J. The role of fixator frame stiffness in the control of fracture healing. An experimental study. J. Biomech. 26, 1027–1035 (1993).

2. Schell, H. et al. The course of bone healing is influenced by the initial shear fixation stability. J. Orthop. Res. 23, 1022–1028 (2005).

3. Betts, D. C. & Müller, R. Mechanical Regulation of Bone Regeneration: Theories, Models, and Experiments. Front. Endocrinol. (Lausanne). 5, (2014).

4. Harris, J. S., Bemenderfer, T. B., Wessel, A. R. & Kacena, M. A. A review of mouse critical size defect models in weight bearing bones. Bone (2013).

5. Claes, L. & Heigele, C. Magnitudes of local stress and strain along bony surfaces predict the course and type of fracture healing. J. Biomech. 32, 255–266 (1999).

6. Prendergast, P., Huiskes, R. & Søballe, K. Biophysical stimuli on cells during tissue differentiation at implant interfaces. J. Biomech. 30, 539–548 (1997).

7. Geris, L., Vander Sloten, J. & Van Oosterwyck, H. In silico biology of bone modelling and remodelling: regeneration. Philos. Trans. Royal Soc. A. 367, 2031–2053 (2009).

8. Lacroix, D. & Prendergast, P. A mechano-regulation model for tissue differentiation during fracture healing: analysis of gap size and loading. J. Biomech. 35, 1163–1171 (2002).

9. Burke, D. P. & Kelly, D. J. Substrate stiffness and oxygen as regulators of stem cell differentiation during skeletal tissue regeneration: a mechanobiological model. PLoS One 7, e40737 (2012).

10. Repp, F., Vetter, A., Duda, G. N. & Weinkamer, R. The connection between cellular mechanoregulation and tissue patterns during bone healing. Med. Biol. Eng. Comput. 53, 829–842 (2015).

11. Vetter, A., Witt, F., Sander, O., Duda, G. & Weinkamer, R. The spatio-temporal arrangement of different tissues during bone healing as a result of simple mechanobiological rules. Biomech. Model. Mechanobiol. 11, 147–160 (2012).

12. Bottlang, M., Mohr, M., Simon, U. & Claes, L. Acquisition of full-field strain distributions on ovine fracture callus cross-sections with electronic speckle pattern interferometry. J. Biomech. 41, 701–705 (2008).

13. Steiner, M., Claes, L., Simon, U., Ignatius, A. & Wehner, T. A computational method for determining tissue material properties in ovine fracture calluses using electronic speckle pattern interferometry and finite element analysis. Med. Eng. Phys. 34, 1521–1525 (2012).

14. Morgan, E. F. et al. Correlations between local strains and tissue phenotypes in an experimental model of skeletal healing. J. Biomech. 43, 2418–2424 (2010).

15. Miller, G. J., Gerstenfeld, L. C. & Morgan, E. F. Mechanical microenvironments and protein expression associated with formation of different skeletal tissues during bone healing. Biomech. Model. Mechanobiol. 14, 1239–1253 (2015).

16. Schulte, F. A., Lambers, F. M., Mueller, T. L., Stauber, M. & Müller, R. Image interpolation allows accurate quantitative bone morphometry in registered micro-computed tomography scans. Comput. Methods Biomech. Biomed. Engin. 17, 539–548 (2014).

17. Schulte, F. A., Lambers, F. M., Kuhn, G. & Müller, R. In vivo micro-computed tomography allows direct three-dimensional quantification of both bone formation and bone resorption parameters using time-lapsed imaging. Bone 48, 433–442 (2011).

18. Webster, D., Schulte, F. A., Lambers, F. M., Kuhn, G. & Müller, R. Strain energy density gradients in bone marrow predict osteoblast and osteoclast activity: a finite element study. J. Biomech. 48, 866–874 (2015).

19. Christen, P. et al. Bone remodelling in humans is load-driven but not lazy. Nat. Commun. 5, 4855 (2014).

20. Christen, P. & Müller, R. In vivo visualisation and quantification of bone resorption and bone formation from time-lapse imaging. Curr. Osteoporos. Rep. 15, 311–317 (2017).

21. Mehta, M., Checa, S., Lienau, J., Hutmacher, D. & Duda, G. N. In vivo tracking of segmental bone defect healing reveals that callus patterning is related to early mechanical stimuli. Eur. Cell. Mater. 24, 71 (2012).

22. Tsitsilonis, S. et al. The effect of traumatic brain injury on bone healing: an experimental study in a novel in vivo animal model. Injury 46, 661–665 (2015).

23. Trüssel, A., Flohr, F., Kuhn, G. A. & Müller, R. Gene expression and local in vivo environment (LivE) imaging of osteocyte subpopulations in trabecular mouse bone. J. Bone. Miner. Res. 30(S1):S100, (2015).

24. Morgan, E. F. et al. Micro-computed tomography assessment of fracture healing: relationships among callus structure, composition, and mechanical function. Bone 44, 335–344 (2009).

25. O’Neill, K. R. et al. Micro-computed tomography assessment of the progression of fracture healing in mice. Bone 50, 1357–1367 (2012).

26. Locher, R. et al. Traumatic brain injury and bone healing: radiographic and biomechanical analyses of bone formation and stability in a combined murine trauma model. J. Musculoskelet. Neuronal. Interact. 15, 309–315 (2015).

27. Gardner, M. J. et al. Differential fracture healing resulting from fixation stiffness variability: a mouse model. J. Orthop. Sci. 16, 298–303 (2011).

28. Weis, J. A. et al. Comparison of microCT and an inverse finite element approach for biomechanical analysis: Results in a mesenchymal stem cell therapeutic system for fracture healing. J. Biomech. (2012).

29. Freeman, T. A., Patel, P., Parvizi, J., Antoci, V. & Shapiro, I. M. Micro-CT analysis with multiple thresholds allows detection of bone formation and resorption during ultrasound-treated fracture healing. J. Orthop. Res. 27, 673–679 (2009).

30. Zwingenberger, S. et al. Establishment of a femoral critical-size bone defect model in immunodeficient mice. J. Surg. Res. 181, e7–e14 (2013).

31. Schell, H. et al. Osteoclastic activity begins early and increases over the course of bone healing. Bone 38, 547–554 (2006).

32. Epari, D. R., Taylor, W. R., Heller, M. O. & Duda, G. N. Mechanical conditions in the initial phase of bone healing. Clin. Biomech. 21, 646–655 (2006).

33. Schulte, F. A. et al. Strain-adaptive in silico modeling of bone adaptation: A computer simulation validated by in vivo micro-computed tomography data. Bone 52, 485–492 (2013).

34. Flaig, C. A highly scalable memory efficient multigrid solver for μ-finite element analyses. (2012).

35. Haffner-Luntzer, M. et al. Evaluation of high-resolution In Vivo MRI for longitudinal analysis of endochondral fracture healing in mice. PLoS One 12, e0174283 (2017).

36. Charles, J. P., Cappellari, O., Spence, A. J., Wells, D. J. & Hutchinson, J. R. Muscle moment arms and sensitivity analysis of a mouse hindlimb musculoskeletal model. J. Anat. (2016).

37. Wehrle, E. et al. Evaluation of longitudinal time-lapsed in vivo micro-CT for monitoring fracture healing in mouse femur defect models. Sci. Rep. 10.1038/s41598-019-53822-x (2019).

38. Christen, P., Ito, K., Galis, F. & van Rietbergen, B. Determination of hip-joint loading patterns of living and extinct mammals using an inverse Wolff’s law approach. Biomech. Model. Mechanobiol. 14, 427–432 (2015).

39. Wehrle, E. et al. In vivo micro-CT based approach for discrimination of physiological and impaired healing patterns in mouse femur defect models. Eur. Cell. Mater. 31, (2016).

40. Kohler, T., Stauber, M., Donahue, L. R. & Müller, R. Automated compartmental analysis for high-throughput skeletal phenotyping in femora of genetic mouse models. Bone 41, 659–67 (2007).

41. Webster, D. J., Morley, P. L., van Lenthe, G. H. & Müller, R. A novel in vivo mouse model for mechanically stimulated bone adaptation-a combined experimental and computational validation study. Comput. Methods Biomech. Biomed. Engin. 11, 435–441 (2008).

42. Mulder, L., Koolstra, J. H., den Toonder, J. M. J. & van Eijden, T. M. G. J. Intratrabecular distribution of tissue stiffness and mineralization in developing trabecular bone. Bone 41, 256–65 (2007).

43. Shefelbine, S. J. et al. Prediction of fracture callus mechanical properties using micro-CT images and voxel-based finite element analysis. Bone 36, 480–488 (2005).

44. Simon, U., Augat, P., Utz, M. & Claes, L. A numerical model of the fracture healing process that describes tissue development and revascularisation. Comput. Methods Biomech. Biomed. Engin. 14, 79–93 (2011).

45. Pistoia, W. et al. Estimation of distal radius failure load with micro-finite element analysis models based on three-dimensional peripheral quantitative computed tomography images. Bone 30, 842–848 (2002).

46. Frost, H. M. Bone “mass” and the “mechanostat”: a proposal. Anat. Rec. 219, 1–9 (1987).

47. Schulte, F. A. et al. Local mechanical stimuli regulate bone formation and resorption in mice at the tissue level. PLoS One 8, e62172 (2013).

