## supplementary material S1 for "The association between mineralised tissue formation and the mechanical local *in vivo* environment: Time-lapsed quantification of a mouse defect healing model"

Institute for Biomechanics

ETH Zurich

Leopold-Ruzicka-Weg 4

8093 Zurich, Switzerland

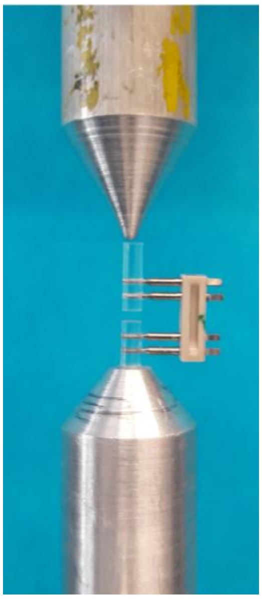

**Supplementary Fig. S1.** Each fixator was assembled and inserted into a PMMA testing rod. The rod was then placed in a custom designed holder on a ZwickRoell II compression tester (ZwickRoell, Ulm, Germany) and compressed quasi-statically to 1 N (preload of 0.05 N) thrice. A 10 N load cell was used (sensitivity of 0.5 %) and the moving platen tip was rounded to prevent non-axial forces and moments upon the specimen. Reported stiffness is the mean of the gradients of the linear force-displacement curve for the three tests in N/mm.

**Supplementary Table S1.** The mean measured stiffness of the PMMA-Fixator constructs.

| Fixator Number | Fixator Stiffness (N/mm) |
|---|---|
| 1 | 23.50 |
| 2 | 23.51 |
| 3 | 23.59 |
| 4 | 23.43 |
| 5 | 23.60 |
| 6 | 23.95 |
| 7 | 22.96 |
| 8 | 23.34 |
| 9 | 22.99 |
| 10 | 24.56 |
| 11 | 25.17 |
| 12 | 25.29 |
| Mean | 23.82 |
| Standard deviation | 0.74 |
